# The impact of eye closure on anticipatory alpha activity in a tactile discrimination task

**DOI:** 10.1101/2021.08.03.454920

**Authors:** Hesham A. ElShafei, Corinne Orlemann, Saskia Haegens

## Abstract

One of the very first observations made regarding alpha oscillations (8–14 Hz), is that they increase in power over posterior areas when awake participants close their eyes. Recent work, especially in the context of (spatial) attention, suggests that alpha activity reflects a mechanism of functional inhibition. However, it remains unclear how eye closure impacts anticipatory alpha modulation observed in attention paradigms, and how this affects subsequent behavioral performance. Here, we recorded magnetoencephalography (MEG) in 33 human participants performing a tactile discrimination task with their eyes open vs. closed. We replicated the hallmarks of previous somatosensory spatial attention studies: alpha lateralization across the somatosensory cortices as well as alpha increase over posterior regions. Furthermore, we found that eye closure leads to (i) reduced task performance, (ii) widespread increase in alpha power, and (iii) reduced anticipatory visual alpha modulation (iv) with no effect on somatosensory alpha lateralization. Regardless of whether participants had their eyes open or closed, increased posterior alpha power and somatosensory alpha lateralization improved their performance. Thus, we provide evidence that eye closure does not alter the impact of anticipatory alpha modulations on behavioral performance. We propose there is an optimal posterior alpha level for somatosensory task performance, which can be achieved through a combination of eye closure and top-down anticipatory attention.

**Significance Statement:** Alpha oscillations are dominant when awake participants have their eyes closed. Furthermore, alpha is known to modulate with anticipatory attention, and has been ascribed a role of active functional inhibition. Surprisingly, the link between anticipatory alpha and eye closure remains unclear. Here we collected MEG data while human participants performed a tactile discrimination task either with their eyes open or closed. Eye closure led to a widespread increase in alpha power, and affected anticipatory visual alpha modulation but not somatosensory alpha lateralization. Importantly, eye closure did not affect the correlation between alpha and task performance. Our findings provide novel insights into how eye closure impacts anticipatory alpha modulation, and how optimal alpha levels for task performance can be achieved differently.

## Introduction

Since the discovery of the cortical alpha rhythm by Hans Berger (1929) almost a century ago, it has been known that a general increase of posterior alpha power occurs when awake participants close their eyes (Adrian & Matthews, 1934). While traditionally the alpha rhythm was associated with a state of cortical idling (Pfurtscheller et al., 1996), more recent work suggests that alpha activity reflects a mechanism of functional inhibition (Foxe & Snyder, 2011; Haegens et al., 2011; Jensen & Mazaheri, 2010; Klimesch et al., 2007). In support of such an inhibitory mechanism, visual spatial attention is known to modulate alpha activity in a lateralized fashion: alpha decreases contralateral to the attended location (Sauseng et al., 2005) and increases contralateral to the ignored location, presumably to suppress distracting input (Kelly et al., 2009; Worden et al., 2000). This lateralized alpha activity correlates with visual detection performance (Händel et al., 2011; Thut et al., 2006). Similar patterns have been observed for the auditory (Banerjee et al., 2011; Frey et al., 2014; Straub et al., 2014; Wöstmann et al., 2016) and somatosensory domains (Anderson & Ding, 2011; Haegens et al., 2011, 2012; Jones et al., 2010).

Importantly, in our previous tactile spatial attention work, we found that somatosensory alpha lateralization was accompanied by an anticipatory increase of posterior alpha power, which positively correlated with tactile discrimination performance. We interpreted this posterior alpha increase to reflect a general inhibition of visual processing to improve tactile performance (Haegens et al., 2010, 2012). An obvious follow-up question is whether a similar posterior alpha increase, and accompanying tactile performance improvement, could be achieved by closing the eyes. Or, in other words, does the anticipatory task-related posterior alpha modulation stem from the same underlying sources as eye-closure related alpha modulation? Another question is how eye-closure induced alpha increase relates to alpha lateralization patterns observed in the context of spatial attention.

Anecdotally, eye closure enhances the concentration on other sensory modalities by suppressing processing of visual input (Glenberg et al., 1998). Eye closure has been shown to boost stimulus responses in somatosensory areas (Brodoehl, Klingner, Stieglitz, et al., 2015; Götz et al., 2017), with mixed findings regarding impact on behavioral performance. To date, the relationship between eye-closure effects and anticipatory alpha modulation has only been investigated in the context of auditory attention: Wöstmann et al. (2020) showed that eye closure increases the general power of alpha oscillations, as well as the modulation of alpha during an auditory attentional task; however, this had no impact on behavioral performance.

Here, we asked whether and how eye-closure induced alpha modulations interact with anticipatory alpha modulations and associated behavioral performance effects. We recorded MEG while participants performed an adapted version of the tactile discrimination task from Haegens et al. (2011), during eyes-open and eyes-closed conditions. First, we asked whether the often-reported eye-closure related power increase extends beyond posterior alpha. Next, we compared the previously reported anticipatory alpha modulations—i.e., somatosensory alpha lateralization and visual alpha increase (Haegens et al., 2012, 2012)—between eye conditions and asked how they interact with the eye-closure related power increase. Finally, we asked whether the relationship between these alpha modulations and task performance differs across eye conditions; specifically, whether visual alpha increase (which we previously interpreted as inhibition of visual processing) is behaviorally relevant in the absence of visual input.

## Materials and Methods

### Participants

Participants were 34 healthy adults (Age: *M* = 25, *SD* = 3.86, range = 20–33 years; 18 female; 30 right handed, 2 left handed, 2 ambidextrous) without neurological or psychiatric disorders, who reported normal hearing and normal or corrected-to-normal vision. The study was approved by the local ethics committee (CMO 2014/288 “Imaging Human Cognition”) and in accordance with the Declaration of Helsinki. Participants gave written informed consent and were remunerated for their participation. One participant was excluded from analysis due to poor data quality.

### Experimental design

Participants performed a tactile discrimination task (Figure 1; task adapted from Haegens et al., 2011) while their brain activity was recorded using MEG. Participants received an electrical stimulus (pulse train of a low or high frequency) to either the right or left thumb. Participants were instructed to determine as fast and accurately as possible whether the perceived stimulus was of low or high frequency, responding via button press with their right index finger (left button press indicated the low frequency; right button press indicated the high frequency). Prior to the stimulus presentation, an auditory cue (verbal “right” or “left”) directed participants’ attention to either their right or left hand. Spatial cues were always valid. Each trial started with a pre-cue interval of 1.2 s followed by the auditory cue (0.2 s), a jittered 1–1.8 s pre-stimulus interval, the tactile stimulus (0.24-s pulse train), a response window of maximum 1.5 s, and finally auditory feedback indicating whether the answer was correct or incorrect.

**Figure 1.**
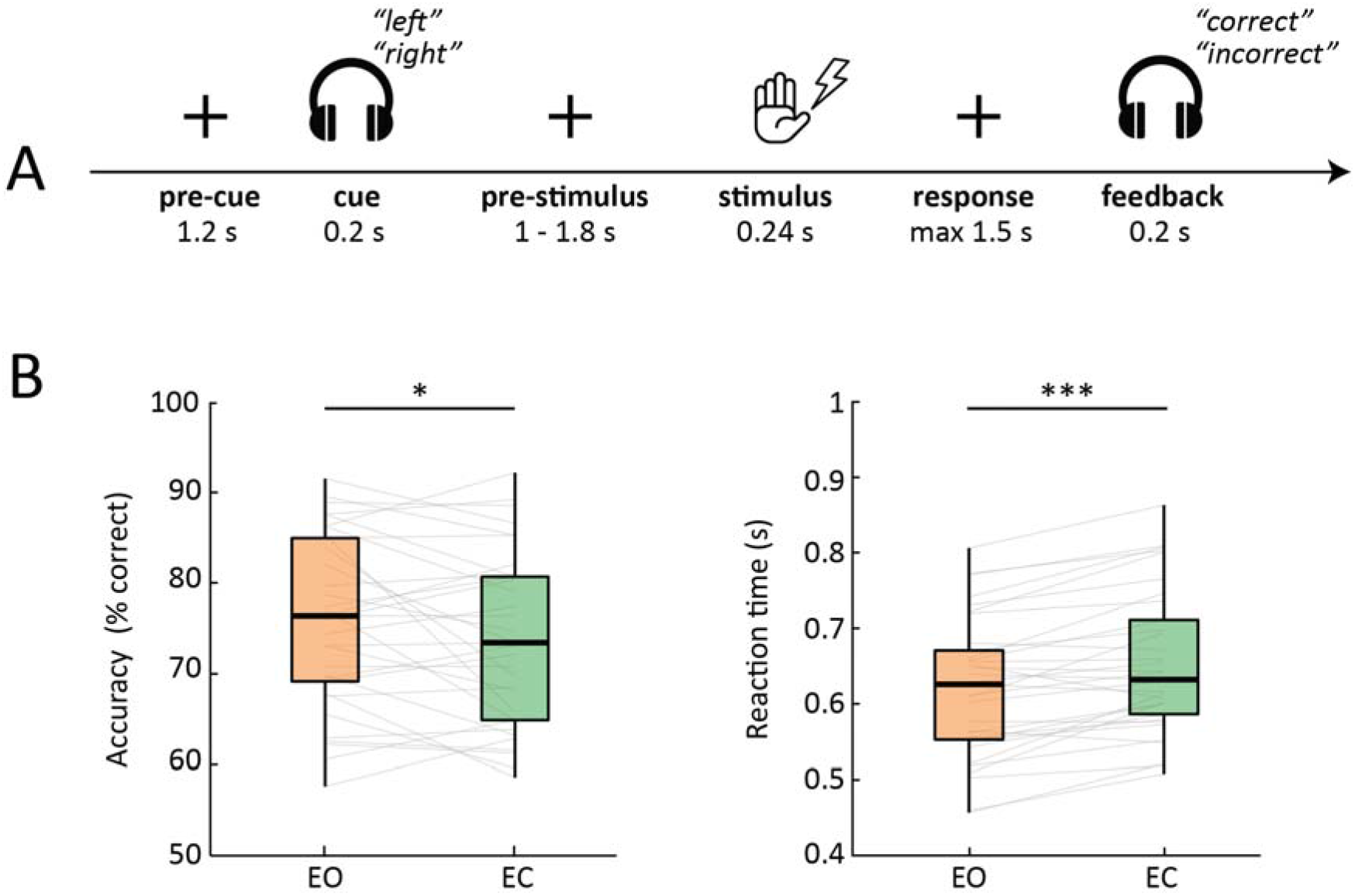
Experimental paradigm and behavioral results. [A] Participants performed a tactile stimulus discrimination task where a 100% valid auditory cue directed attention either to their right or left hand in an eyes-open (EO) and an eyes-closed (EC) condition. Participants had to discriminate between two target frequencies, presented as electrical pulse trains to the cued thumb. [B] Accuracy (left panel) and reaction time (right) for the EO and EC conditions. Behavioral performance was significantly worse when participants had their eyes closed both in terms of lower accuracy and slower RT. *p<0.05; **p<.01; ***p<.001.

Participants performed this task under two conditions: an eyes-open (EO) and an eyes-closed condition (EC). Conditions were presented in a counter-balanced block-design of four blocks per condition with 76 trials each, resulting in a total of 304 trials per condition. During the EO condition, participants were instructed to fixate on a fixation cross in the middle of the screen. For the EC condition, participants kept their eyes closed for the duration of the block. After each block, participants were presented with a short questionnaire to rate their sleepiness level (very sleepy, sleepy, awake, very awake). Prior to the experiment, participants performed four training blocks (two per condition, 12 trials per block), during which they were familiarized with the task.

### Stimulus presentation

We used the same setup as in Haegens et al. (2011): Electrical stimuli were delivered with two constant-current high-voltage stimulators (Digitimer Ltd, Model DS7A) to the right and left thumb. The intensity (*M*_right_ = 6.4 mA, range = 3.9–9.5 mA; *M*_left_ = 5.5 mA, range = 3.2–9.9 mA) of the 0.2-ms electric pulses was set to 150% of the participant’s sensory threshold level. This level was established during a practice session before the recordings, for each thumb independently. Low (either 25 or 33.3 Hz) and high frequencies (41.7, 50, or 66.7 Hz) were determined for each participant individually to ensure successful execution of the task, above chance level but below ceiling performance. Auditory cues and feedback (0.2-s length each) were computer-generated and presented binaurally through air-conducting tubes.

### Data acquisition

Whole-head MEG data were acquired at a sampling frequency of 1200 Hz with a 275-channel MEG system with axial gradiometers (CTF MEG Systems, VSM MedTech Ltd.) in a magnetically shielded room. Six permanently faulty channels were disabled during the recordings, leaving 269 recorded MEG channels. Three fiducial coils were placed at the participant’s nasion and both ear canals, to provide online monitoring of participant’s head position (Stolk et al., 2013) and offline anatomical landmarks for co-registration. Eye position was recorded using an eye tracker (EyeLink, SR Research Ltd.). Upon completion of the MEG session, participant’s head shape and the location of the three fiducial coils were digitized using a Polhemus 3D tracking device (Polhemus, Colchester, Vermont, United States). Anatomical T1-weighted MRIs were obtained during a separate session. To improve co-registration of the MRIs and MEG data, earplugs with a drop of Vitamin E were placed at participant’s ear canals during MRI acquisition. These anatomical scans were used for source reconstruction of the MEG signal.

### Pre-processing

MEG data were preprocessed offline and analyzed using the FieldTrip toolbox (Oostenveld et al., 2011) and custom-built MATLAB scripts. The MEG signal was epoched based on the onset of the somatosensory stimulus (t= −4 to 3 s). The data were downsampled to a sampling frequency of 300 Hz, after applying a notch filter to remove line noise and harmonics (50, 100, and 150 Hz). Bad channels and trials were rejected via visual inspection before independent component analysis (Jung et al., 2001) was applied. Subsequently, components representing eye-related and heart-related artefacts were projected out of the data (on average, eight components were removed per participant). Finally, for the resulting data, outlier trials of extreme variance were removed. This resulted in an average of 537 (± 7 SEM) trials and 268 channels per participant for the reported analyses.

### Spectral analysis

First, we calculated the planar representation of the MEG field distribution from the single-trial data using the nearest-neighbor method. This transformation makes interpretation of the sensor-level data easier as the signal amplitude is typically maximal above a source. Next, we computed spectral representations for two 1-s time windows: the pre-stimulus window and the pre-cue window, aligned to stimulus and cue onset, respectively. Each window was multiplied with a Hanning taper, and power spectra (1–30 Hz; 1-Hz resolution) were computed using a fast Fourier transform (FFT) approach. Additionally, for a time-resolved-representation of the spectral power distribution, we computed time-frequency representations (TFRs) of the power spectra for the full trials per experimental condition. To this end we used an adaptive sliding time window of five cycles length per frequency (Δt = 5/f; 20-ms step size).

### Alpha peak frequency

In order to investigate how eye closure impacts alpha activity we computed the individual alpha peak frequencies for each participant, separately for occipital and centroparietal sensor-level regions of interest (ROIs), and separately for the EO and EC conditions. We determined participants’ peak frequencies within a broad alpha range (7–14 Hz) during the pre-stimulus interval (−1 to 0 s). As intra-individual alpha peaks did not significantly vary with condition (F(1, 32) = 0.46, p = 0.5, ANOVA) or ROI (F(1, 32) = 1.04, p = 0.31), nor their interaction (F(1, 32) = 0.17, p = 0.67), we computed one average peak for each participant (*M* = 10 Hz, range = 7–13 Hz). Using individual alpha peak frequency allows taking into account inter-individual variability, and provides a more accurate estimation of alpha activity than when using a fixed frequency band (Haegens et al., 2014). All further analysis was computed using these individual alpha peaks, with spectral bandwidth of ±1 Hz, unless indicated otherwise.

### Statistical analysis

In order to investigate whether power differences between the EO and the EC conditions were significant, we used nonparametric cluster-based permutation tests (Maris & Oostenveld, 2007). In brief, this test first calculates paired t-tests for each sensor at each time and/or frequency point, which are then thresholded at p < 0.05 and clustered on the basis of spatial, temporal, and/or spectral adjacency. The sum of t-values within each cluster is retained, and the procedure is repeated 1000 times on permuted data in which the condition assignment within each individual is randomized. On each permutation, the maximum sum is retained. Across all permutations, this yields a distribution of 1000 maximum cluster values. From this distribution, the probability of each empirically observed cluster statistic can be derived (evaluated at alpha = 0.05).

We used this permutation test to investigate the impact of eye closure on (i) global oscillatory power, by contrasting power in the pre-stimulus interval between eye conditions, (ii) anticipatory visual alpha activity, by contrasting pre-stimulus baseline-normalized power between eye conditions, for each cue separately, and (iii) somatosensory alpha activity, by contrasting the pre-stimulus attention modulation index, calculated as (attention-left - attention-right) / (attention-left + attention-right) between eye conditions.

In order to investigate the impact of pre-stimulus alpha activity on behavioral performance, we focused our analysis on visual and somatosensory ROIs that were defined in sensor space. For the somatosensory ROIs, our selection was data-based, i.e., per hemisphere we selected 10 sensors with the maximum evoked response to contralateral tactile stimulation. For the visual ROIs, as our design lacked visual stimuli, our selection included 10 left and 10 right occipital sensors. One participant was excluded from analysis due to poor data quality. Note that for alpha power in the visual ROIs, we use the term “absolute” modulation to denote overall non-baseline-normalized power in the pre-stimulus window, while the term “anticipatory” denotes the baseline-normalized power in the same pre-stimulus window.

### Alpha lateralization index

To capture the relative pre-stimulus somatosensory alpha distribution over both hemispheres in one measure, we computed a lateralization index of alpha power (Haegens et al., 2011; Thut et al., 2006) for each participant, using individual somatosensory ROIs: alpha lateralization index = (alpha-ipsilateral - alpha-contralateral) / (alpha-ipsilateral + alpha-contralateral). This index gives positive values if alpha power is higher over the ipsilateral hemisphere and/or lower over the contralateral hemisphere (with contra- and ipsilateral sides defined with respect to the spatial cue). Negative values arise if alpha power activity is lower over the ipsilateral hemisphere and/or higher over the contralateral hemisphere.

### Source reconstruction

In order to localize the generators of the sensor-level spectrotemporal effects, we applied the frequency-domain adaptive spatial filtering technique of dynamical imaging of coherent sources (Gross et al., 2001). For each participant, an anatomically realistic single-shell headmodel based on individual T-1 weighted anatomical images was generated (Nolte, 2003). The brain volume of each individual subject was divided into a grid with a 0.5-cm resolution and normalized toward a template MNI brain using non-linear transformation. For each grid point, leadfields were computed with a reduced rank, which removes the sensitivity to the direction perpendicular to the surface of the volume conduction model. This procedure ensures that each grid-point represents the same anatomical location across all participants by taking into account the between-subject difference in brain anatomy and head shape.

Data from all conditions of interest were concatenated in order to compute the cross-spectral density (CSD) matrices (multitaper method (Mitra & Pesaran, 1999)). Leadfields for all grid points along with the CSD matrices were used to compute a common spatial filter (i.e., common for all trials and conditions) that was used to estimate the spatial distribution of power for time-frequency windows of interest highlighted in the previous analysis. The source orientation was fixed to the dipole direction with the highest strength.

## Results

### Eye closure impairs performance

Performance over all 33 participants for both eye conditions combined was an average accuracy of 74.4% (*SD* = 9.96%) and an average reaction time (correct trials only) of 0.64 s (*SD* = 0.1 s). Participants were more accurate (t(32) = 2.32, p = 0.023, paired-test, mean EO = 75.7% + 9.9 SD, mean EC = 73.7% ± 9.9 SD) and faster (t(32) = −6.8, p < 0.001, mean EO = 0.62 s ± 0.1 SD, mean EC = 0.65 s ± 0.1 SD) at discriminating the frequency of the tactile stimuli in the EO condition in comparison to the EC condition (Figure 1B).

Further, we investigated the impact of eye closure (two levels: EC and EO) and block order (four levels: first, second, third and fourth) on the sleepiness score reported at the end of each block. We found a main effect of eye condition (F(1,26) = 9.7, p = 0.004, ANOVA), with participants reporting being more awake when they had their eyes open. In addition, we found a main effect of block order (F(3,78) = 5.32, p = 0.009), with participants reporting being more awake in the first block in comparison to the second (t(26) = −3.15, p = 0.014, posthoc paired t-test), third (t(26) = −3.45 p = 0.005) and fourth (t(26) = −3.15, p = 0.014), with no significant interaction (F(3,78) = 1.11, p = 0.35). Note that differences in sleepiness scores did not correlate with differences in behavioral performance between eye conditions (RT: r(26) = −0.19, p = 0.32; accuracy: r(26) = 0.22, p = 0.25).

### Eye closure boosts global oscillatory activity

In order to investigate the impact of eye closure on overall oscillatory power, we contrasted power spectra (1–30 Hz) during the pre-stimulus window between the EO and the EC conditions (Figure 2). We found that power was higher for EC than EO (cluster-corrected p < 0.001), both in the alpha (6–12 Hz) and in the beta range (17–30 Hz). The alpha cluster was widespread with a spectral peak at 10 Hz, while the beta cluster was concentrated towards posterior sensors, showing the highest difference between conditions around 20 Hz. While in this study we focused on alpha activity, as a control we compared event-related fields (ERFs) between eye conditions and found no differences (cluster-corrected p > 0.5).

**Figure 2.**
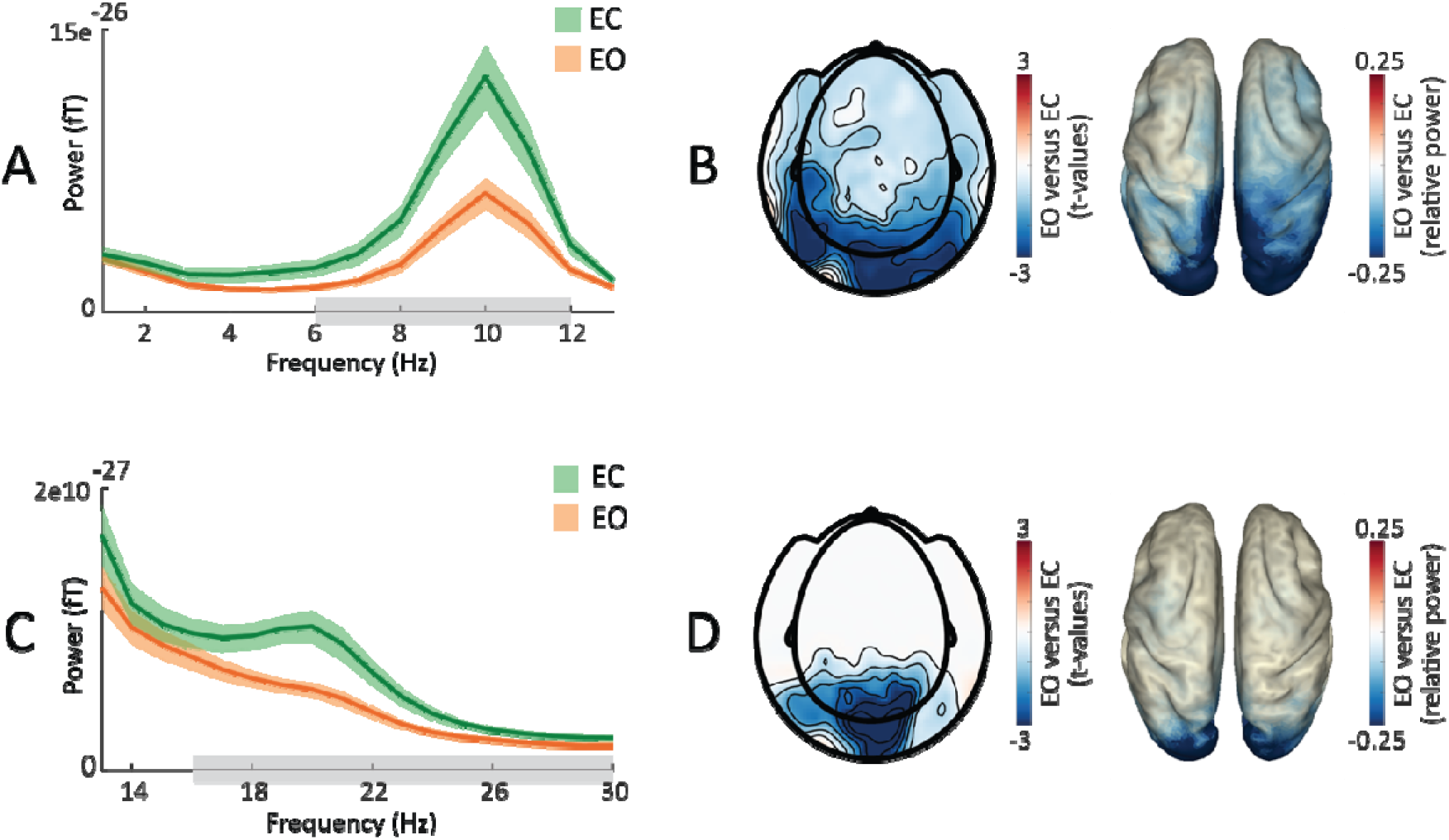
Impact of eye closure on global power. [A] Average absolute occipital power (1–13 Hz) during the pre-stimulus window (t = −1 to 0 s) for the EC (green) and EO (orange) conditions (shading reflects between-participant SEM). Alpha power was significantly higher in the EC condition compared to the EO condition. Grey bars indicate significant differences between conditions. [B] Topography of significant (masked at p < 0.05) cluster t-values for the alpha band for EO vs. EC (as marked in A) on sensor level (left panel) and power distribution of these differences in source space (right). [C] Same as panel A for 13–30 Hz. Beta power was significantly higher in the EC condition compared to the EO condition. [D] Same as panel B for the beta band (as marked in C).

### Eye closure impacts anticipatory visual alpha modulation

In order to investigate the impact of eye closure on anticipatory alpha modulation, we first contrasted alpha power between the pre-stimulus and the baseline windows. We found a pre-stimulus decrease of alpha power over left central sensors vs. baseline, for both EO and EC conditions (Figure 3AB; cluster-corrected p = 0.005). Furthermore, we observed a pre-stimulus increase of posterior alpha power (p = 0.001), which was exclusive to the EO condition. Next, we directly contrasted the baseline-normalized pre-stimulus alpha between EO and EC conditions, separately for each attention condition (i.e., attend left and right). For both attention conditions, we found higher posterior alpha power in the EO condition compared to the EC condition (cluster-corrected p < 0.001; Figure 3CD). This result reflects an increase of visual alpha power during the pre-stimulus interval vs. baseline in the EO condition, an effect that was absent in the EC condition. Hence, despite an overall increase of alpha power with eye closure, the anticipatory posterior alpha modulation during the pre-stimulus interval was higher for open eyes.

**Figure 3.**
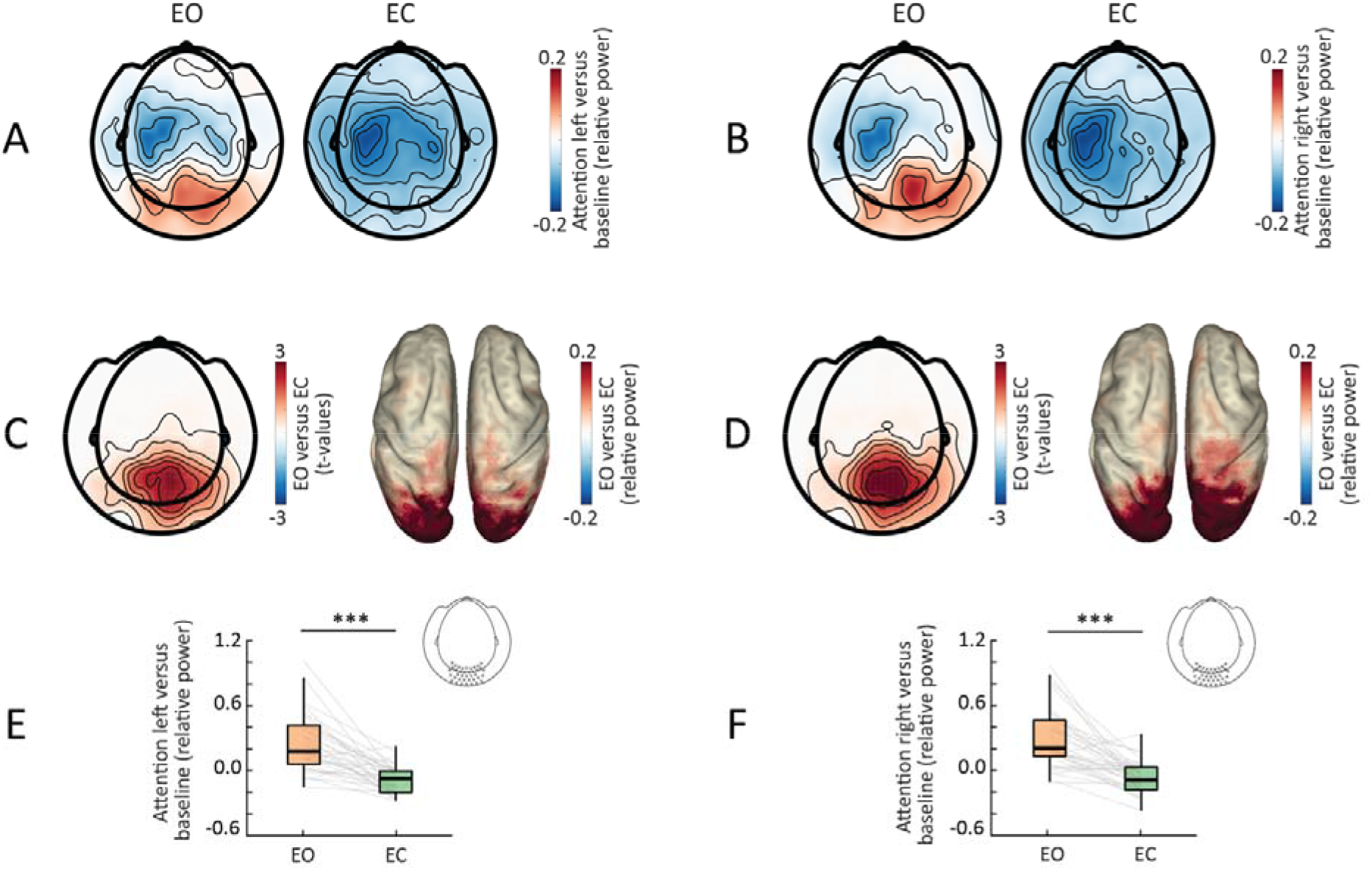
Impact of eye closure on anticipatory visual alpha modulation. [A] Topography of the normalized pre-stimulus alpha power modulation for the attention-left condition (i.e., pre-stimulus window vs. baseline) for EO (left panel) and EC (right). [B] Same as A for the attention-right condition. [C] Topography of significant (masked at p < 0.05) cluster t-values for EO vs. EC for the attention-left condition on sensor level (left panel), and power distribution of these differences in source space (right). [D] Same as C for the attention-right condition. [E] Normalized occipital pre-stimulus alpha power for the attention-left condition (included sensors marked in topography inset), showing significant difference between eye conditions. [F] Same as E for the attention-right condition. *p<0.05; **p<.01; ***p<.001.

### Eye-closure related and anticipatory alpha modulations are spatially distinct

To address the question of whether eye-closure induced modulations and anticipatory alpha modulations share the same underlying cortical generators (i.e., localize to the same cortical regions), we compared the maxima of these effects in source space. For each participant, we identified the voxel displaying the maximal difference in absolute alpha power in the EO and the EC conditions, and the voxel displaying the maximal anticipatory pre-stimulus alpha power modulation. We then contrasted the x- y- and z- coordinates of these maxima using paired t-tests. We found that maxima differed in their distribution along the y-axis (t(32) = −2.83, p = 0.007 paired t-test) and the z-axis (t(32) = −3.7, p < 0.001). In other words, maxima of the anticipatory alpha modulations were located more anterior and superior in comparison to the eye-closure induced modulations (Figure 4), with no differences in the distribution along the x-axis (i.e., left vs. right; t(32) = 0.36, p = 0.71).

**Figure 4.**
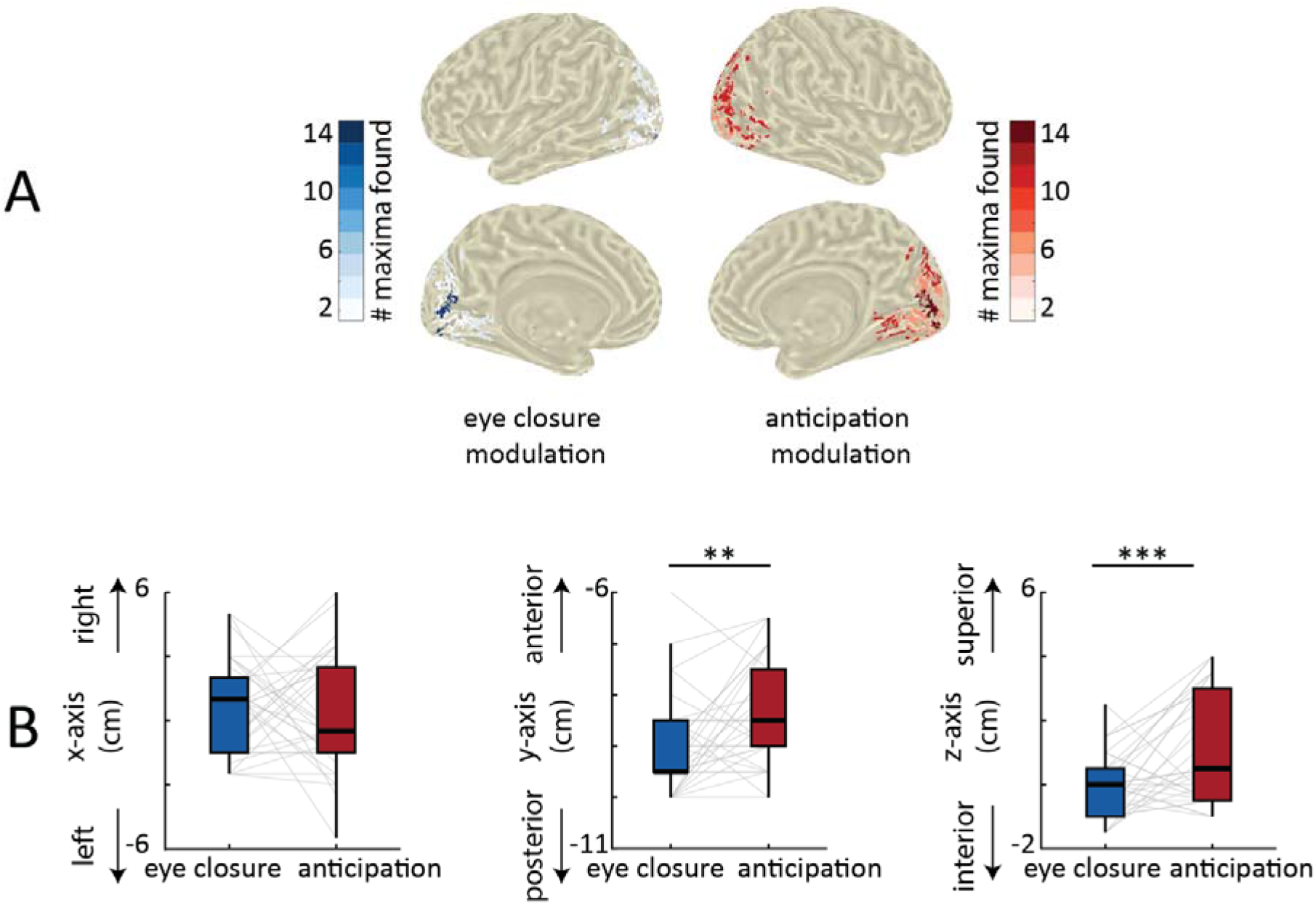
Localization differences between eye-closure and anticipatory alpha modulations. [A] Distribution of the eye-closure (in blue, left) and anticipatory (in red, right) alpha modulations in source space. For visualization purposes, maximas from each modulation were transposed on one hemisphere. [B] Topography of significant (masked at p < 0.05) cluster t-values for eye closure vs. anticipatory alpha modulations. [C] Maxima coordinates along the x-axis (left), y-axis (middle) and z-axis (right). *p<0.05; **p<.01; ***p<.001.

### Eye closure does not impact somatosensory alpha modulation

In order to investigate how eye closure impacts anticipatory somatosensory alpha modulation, we contrasted the pre-stimulus attention modulation index (i.e., attention left vs. right) between EO and EC conditions. While there was a significant attention modulation—i.e., a pattern of lateralized sensorimotor alpha power (left increase p = 0.007; right decrease p < 0.001) when contrasting left vs. right attention conditions—no significant differences were found between eye conditions (p = 0.34; Figure 5). Thus, while both overall and anticipatory visual alpha activity differed between eye conditions, anticipatory somatosensory alpha modulation was not affected by eye closure.

**Figure 5.**
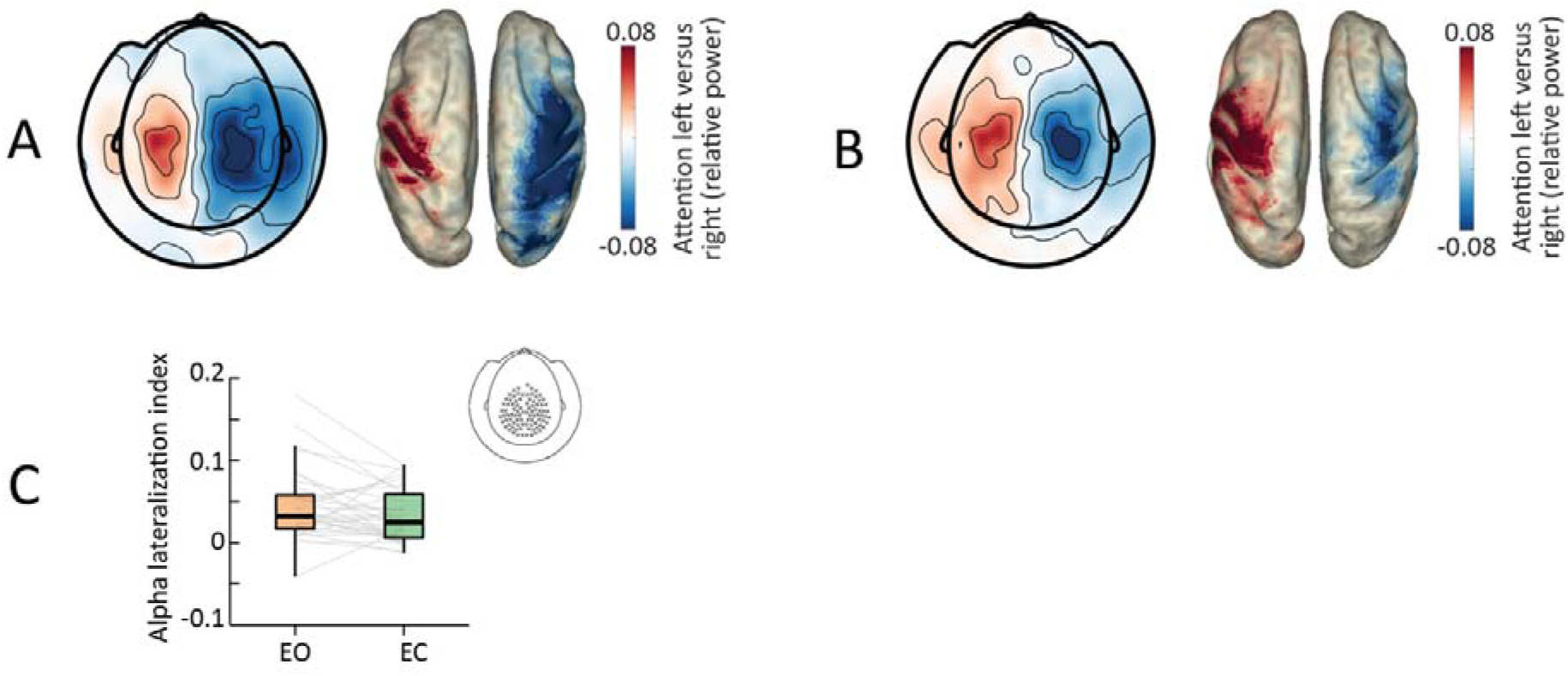
Impact of eye closure on somatosensory alpha lateralization. [A] Topography of the attention-left vs attention-right anticipatory alpha power modulation for the EO condition (left panel), and power distribution of this modulation in source space (right). This modulation localizes to somatomotor regions with higher alpha power in ipsilateral and lower alpha power in contralateral regions. [B] TFRs showing the lateralized power modulation in the EO condition. Left-hemispheric sensors were mirrored to combine them with the right-hemispheric sensors. [C] Same as A for the EC condition. [D] Same as B for the EC condition. [E] Pre-stimulus alpha lateralization index (included sensors marked in topography inset), showing no significant difference between eye conditions.

### Eye closure does not impact the link between anticipatory alpha and behavioral performance

Finally, we investigated the impact of eye closure on the link between pre-stimulus alpha modulation and behavioral performance. First, we analyzed the relationship between pre-stimulus visual alpha power, both absolute (non-baseline normalized) and anticipatory (baseline-normalized) modulations, and performance, by binning the data based on correct vs. incorrect responses, and fast vs. slow RTs (divided by a median split).

For absolute visual alpha power and accuracy (Figure 6A), we found a significant main effect of accuracy (F (1, 31) = 15.2, p < 0.001, ANOVA) with absolute visual alpha power being higher in correct trials in comparison to incorrect trials. In addition, we found a significant main effect of eye condition (F(1, 31) = 26.92, p < 0.001) and no significant interaction between eye condition and accuracy (F(1, 31) = 1.15, p = 0.29). For absolute visual alpha power and RT (Figure 6B), we found a significant main effect of RT (F(1, 31) = 6.11, p = 0.02, ANOVA) with absolute visual alpha power being higher in fast trials in comparison to slow trials. In addition, we found a significant main effect of eye condition (F(1, 31) = 31.53, p < 0.001) and no significant interaction between eye condition and RT (F(1, 31) = 0.65, p = 0.42). In sum, absolute visual alpha power predicted more accurate and faster responses, regardless of eye condition.

**Figure 6.**
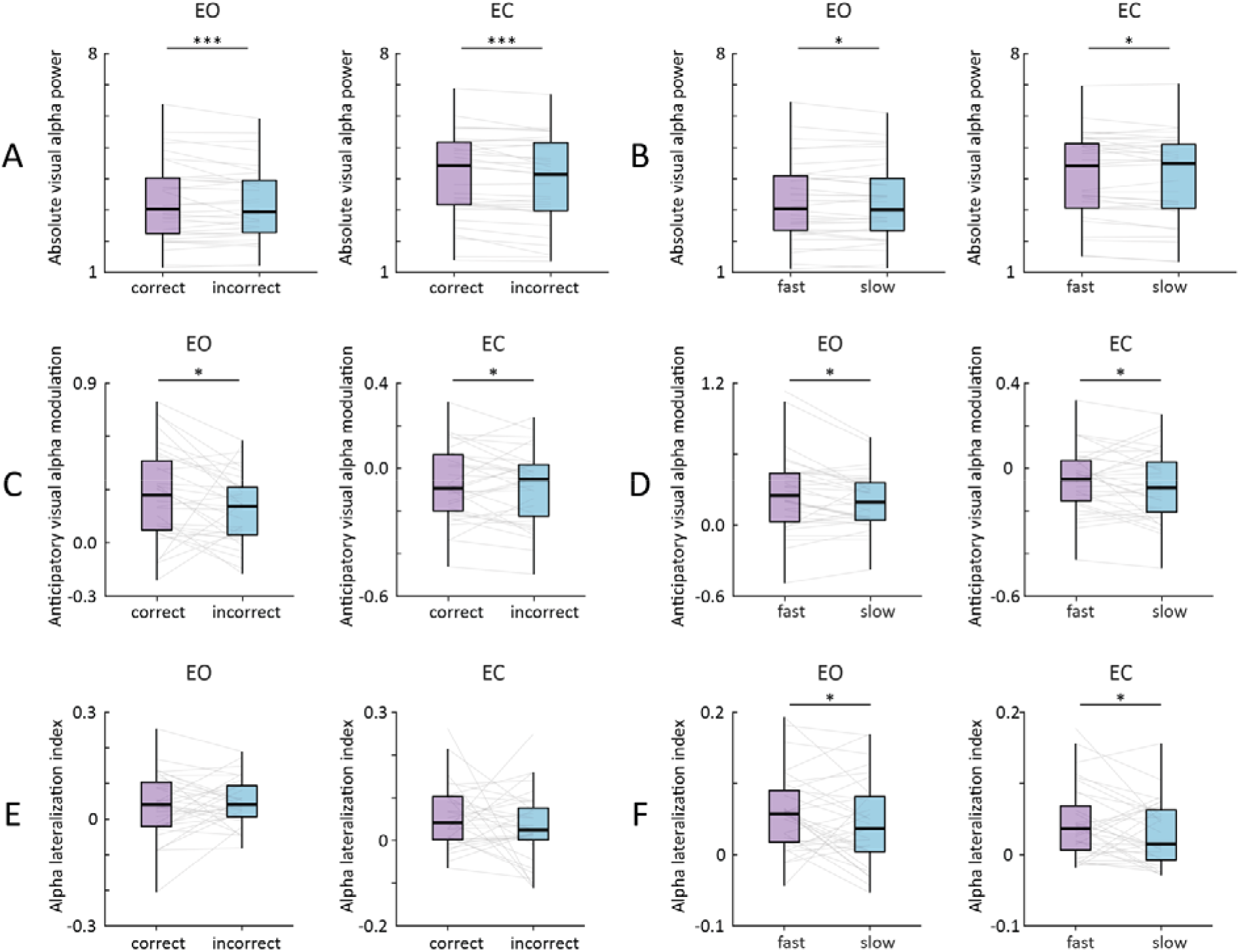
Impact of eye closure on the relationship between alpha and performance. [A] Absolute (non-baseline corrected) pre-stimulus visual alpha power in EO (left panel) and EC (right panel) conditions for correct vs. incorrect trials. Absolute visual alpha power was higher for correct trials, regardless of eye condition. [B] Same as A for fast vs. slow trials. Absolute visual alpha power was higher for fast trials, regardless of eye condition. [C] Same as A for anticipatory visual alpha modulation (baseline corrected) in EO (left panel) and EC (right panel) conditions for correct vs. incorrect trials. Anticipatory visual alpha power was higher for correct trials, regardless of eye condition. [D] Same as C for fast vs. slow trials. Anticipatory visual alpha power was higher for fast trials, regardless of eye condition. [E] Same as C for somatosensory alpha lateralization index. No significant differences were found between conditions. [F] Same as E for fast vs. slow trials. Somatosensory alpha lateralization was higher for fast trials, regardless of eye condition. *p<0.05; **p<.01; ***p<.001.

For anticipatory visual alpha power and accuracy (Figure 6C), we found a significant main effect of accuracy (F (1, 31) = 4.84, p = 0.035, ANOVA) with anticipatory visual alpha power being higher in correct trials in comparison to incorrect trials. In addition, we found a significant main effect of eye condition (F(1, 31) = 69.88, p < 0.001) and no significant interaction between eye condition and accuracy (F(1, 31) = 1.77, p = 0.19). For anticipatory visual alpha power and RT (Figure 6D), we found a significant main effect of RT (F(1, 31) = 7.39, p = 0.01, ANOVA) with anticipatory visual alpha power being higher in fast trials in comparison to slow trials. In addition, we found a significant main effect of eye condition (F(1, 31) = 41.21, p < 0.001) and no significant interaction between eye condition and RT (F(1, 31) = 1.04, p = 0.31). In sum, anticipatory visual alpha modulation predicted more accurate and faster responses, regardless of eye condition.

For somatosensory alpha lateralization and accuracy (Figure 6E), we did not find a significant main effect of accuracy (F (1, 31) = 0.39, p = 0.53, ANOVA) nor a significant main effect of eye condition (F (1, 31) = 0.001, p = 0.98), nor a significant interaction between eye condition and accuracy (F (1, 31) = 1.19, p = 0.28). For somatosensory alpha lateralization and RT (Figure 6F), we found a significant main effect of RT (F (1, 31) = 5.31, p = 0.027, ANOVA) with somatosensory alpha lateralization being higher for faster trials. We found neither a significant main effect of eye condition (F (1, 31) = 2.47, p = 0.12) nor a significant interaction between eye condition and RT (F (1, 31) = 0.001, p = 0.98). In sum, somatosensory alpha lateralization predicted faster responses, regardless of eye condition.

## Discussion

In a follow-up on our previous work (Haegens et al., 2010, 2011, 2012), we investigated how eye-closure related alpha modulations interact with anticipatory alpha dynamics and subsequent behavioral performance during a tactile spatial attention task. We found that task performance was reduced with eye closure. While eye closure led to a widespread increase in alpha power, this only affected anticipatory visual alpha modulation, with somatosensory alpha lateralization being the same across eyes-open and -closed conditions. Regardless of whether participants had their eyes open or closed, increases in posterior alpha power and somatosensory alpha lateralization improved their performance.

### Eye closure impacts global state

Participants were less accurate and slower to discriminate tactile stimuli when their eyes were closed. While there have been several reports of a positive impact of eye closure on performance (e.g., perceptual sensitivity: Brodoehl, Klingner, Stieglitz, et al., 2015; memory retrieval: Parker & Dagnall, 2020; Vredeveldt et al., 2011), other studies have reported no effects (e.g., memory retrieval: Bastarrika-Iriarte & Caballero-Gaudes, 2019; selective attention: Wöstmann et al., 2020) or negative impact (somatosensory discrimination Götz et al., 2017). Differences in paradigms (attention versus memory) and sensory modalities (auditory versus somatosensory) between these various reports renders it difficult to define common factors that govern the interaction between eye closure and behavioral performance. Nevertheless, Götz et al. (2017) argue that for tactile perception, eye closure might boost sensitivity but hinder discriminability, possibly due to the dependence of tactile discriminability upon extrastriate visual processing (Sathian & Zangaladze, 2002). Following this logic, in our tactile discrimination task eye closure diminishes extrastriate visual processing, leading to worse behavioral performance.

Simultaneous with this behavioral deterioration, and as has been long known (e.g., Adrian & Matthews, 1934; Geller et al., 2014; Wöstmann et al., 2020), alpha power increased with eye closure. This increase was widespread, extending beyond occipital regions, and additionally included frequency ranges neighboring the alpha band (i.e., theta and beta). This observation supports the idea that eye closure does not only reflect a disengagement of visual areas, but rather a cortical state transition (Barry et al., 2007; Harris & Thiele, 2011; Marx et al., 2004). One interesting question is whether the observed oscillatory shifts are dependent on (lack of) light input or eye closure per se. Findings from resting state studies have been contradictory, with reports that alpha power is modulated by light input but not eye closure itself, and vice versa (Ben-Simon et al., 2013; Jao et al., 2013). Future research should investigate how light input impacts the interaction between eye closure and oscillatory dynamics during active tasks.

### Eye closure versus anticipatory attention

Although eye closure led to a general increase of alpha power, we found a significant reduction of *anticipatory* visual alpha modulation in comparison to the eyes-open condition, with the maxima of this latter phenomenon extending more anterior than the global alpha increase. Somatosensory alpha lateralization was not affected by eye closure. These observed alpha modulations are in line with the proposal that alpha power reflects a functional mechanism of inhibition (Foxe & Snyder, 2011; Haegens et al., 2011; Jensen & Mazaheri, 2010; Klimesch et al., 2007) that regulates cortical excitability to gate information from task-irrelevant regions (here: visual and ipsilateral somatosensory cortices) to task-relevant ones (contralateral somatosensory cortex).

To our knowledge, only two previous studies investigated the interaction between eye-closure induced and task-related alpha modulations. Both studies, using auditory paradigms without a spatial component, reported an eye-closure related increase in alpha power (Bastarrika-Iriarte & Caballero-Gaudes, 2019; Wöstmann et al., 2020). Wöstmann et al. (2020) found that eye closure enhances the attentional modulation of alpha power, and Bastarrika-Iriarte & Caballero-Gaudes (2019) found that eye closure enhances the event-related alpha power increase. Neither study found an effect of eye closure on performance (i.e., accuracy). In their study, Wöstmann et al. (2020) presented to-be-attended and to-be-ignored speech streams binaurally, i.e., attention was equally distributed across auditory cortices. Importantly, they found that eye closure enhances attentional modulation primarily in non-auditory (task-irrelevant) parieto-occipital regions. This mirrors our finding that eye closure only impacts anticipatory visual (task-irrelevant) alpha modulation. Note that since somatosensory demands are equivalent across eye conditions, and any non-lateralized effects are subtracted out in our lateralization index, it follows that anticipatory somatosensory alpha remains unaffected by eye closure.

We found that both absolute and anticipatory visual alpha increase were associated with faster and more accurate responses in both eye conditions. This aligns with our previous findings in the somatosensory (Haegens et al., 2010, 2012) and the auditory domains (e.g., ElShafei et al., 2018), demonstrating that in non-visual tasks, visual alpha increase facilitates behavioral performance. In addition, we found that anticipatory somatosensory lateralization was associated with faster responses, regardless of eye condition. The absence of an effect of somatosensory lateralization on accuracy contradicts our previous findings that lateralization leads to better accuracy (Haegens et al., 2011; Haegens et al., 2012). However, a key difference with our current study is the presence of distracting (competing) tactile stimuli in our previous work. If alpha controls inhibition, it is conceivable that the link between somatosensory lateralization and accuracy is to a degree dependent on the presence of distracting somatosensory stimuli that require suppressing, and we may therefore not have been as sensitive to such effects here.

Critically, all observed alpha-performance correlations were independent of eye-closure condition; i.e., eye closure did not impact the relationship between alpha dynamics and behavioral performance. Furthermore, both global and anticipatory visual alpha changes showed similar relationships with task performance, suggesting a general (functional inhibitory) role for alpha, regardless of driving/modulatory factor behind the observed alpha dynamics. We propose that posterior alpha reflects the inhibition of task-irrelevant visual processing, and that in the presence of visual input (eyes-open condition) an increase in visual alpha power is required to achieve this, while in the absence of visual input (eyes-closed condition), visual alpha power is already elevated, hence reducing the need for additional anticipatory modulation (Figure 7).

**Figure 7.**
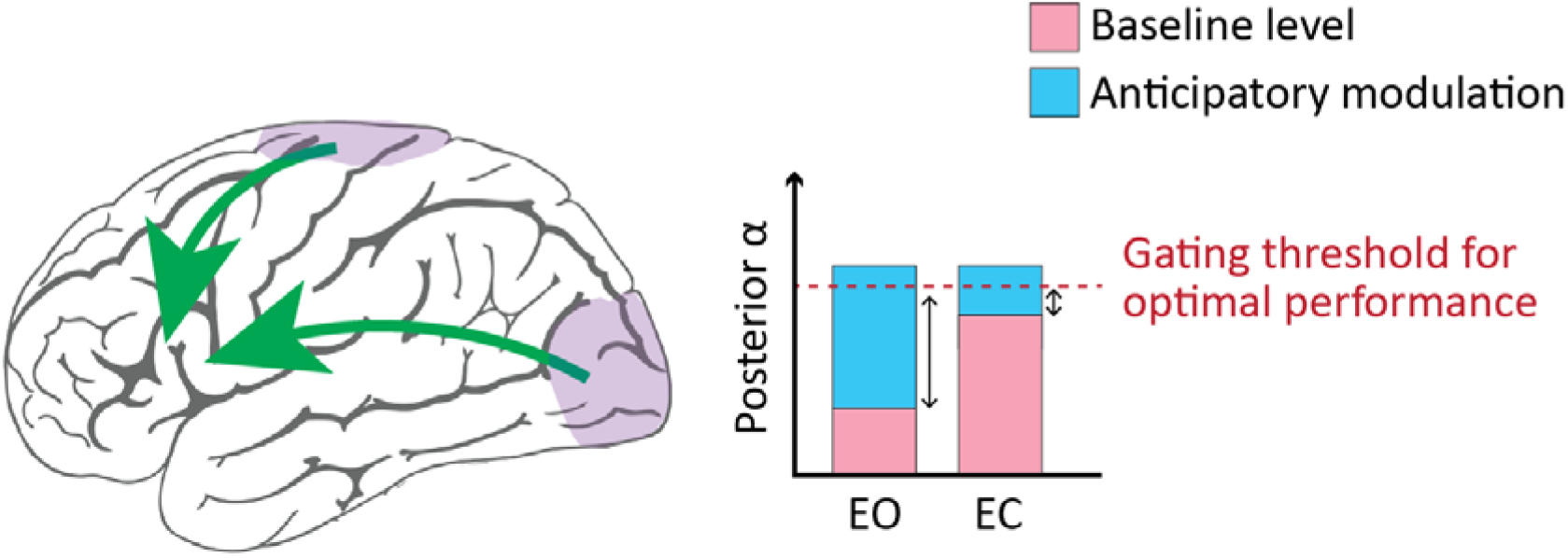
Information gating and eye closure. In the EO baseline interval, information processing is equivalent across task-relevant somatosensory and task-irrelevant visual regions. Thus, in the pre-stimulus interval anticipatory modulation drives alpha levels to the optimal gating threshold at which information flow is gated away from visual regions by inhibiting the processing of visual input. In the EC baseline interval information processing is already diminished due to the absence of visual input. However, alpha level has not yet reached the optimal threshold to entirely gate information flow. Thus, in the pre-stimulus interval, alpha level is further heightened to reach the gating threshold and thus inhibiting information processing in visual regions.

## Conclusion

The present study dissociates for the first time eye-closure induced alpha and anticipatory alpha modulations in the somatosensory domain. We demonstrate that while eye closure boosts global alpha power, it dampens anticipatory visual alpha modulation with no impact on somatosensory lateralization. Finally, we show that eye closure does not alter the impact of alpha dynamics on behavioral performance. Combined, this suggests there is an optimal posterior alpha level for somatosensory task performance, which can be achieved both through eye closure and top-down anticipatory attention. Our findings provide further support for a general inhibitory or gating role for the alpha rhythm.

## Notes

**Conflict of interest statement**, The authors declare that the research was conducted in the absence of any commercial or financial relationships that could be construed as a potential conflict of interest.

### Competing Interest Statement

The authors have declared no competing interest.

